# Zinc finger protein Zfp335 controls early T cell development and survival through β-selection-dependent and -independent mechanisms

**DOI:** 10.1101/2021.11.21.469458

**Authors:** Xin Wang, Anjun Jiao, Lina Sun, Wenhua Li, Biao Yang, Yanhong Su, Renyi Ding, Cangang Zhang, Haiyan Liu, Xiaofeng Yang, Chenming Sun, Baojun Zhang

## Abstract

T cell development in the thymus undergoes the process of differentiation, selective proliferation and survival from CD4^-^CD8^-^ double negative (DN) stage to CD4^+^CD8^+^ double positive (DP) stage prior to the formation of CD4^+^ helper and CD8^+^ cytolytic T cells ready for circulation. Each developmental stage is tightly regulated by sequentially-operating molecular networks, of which only limited numbers of transcription regulators have been deciphered. Here we identified Zfp335 transcription factor as a new player in the regulatory network controlling thymocyte development. We demonstrate that Zfp335 intrinsically controls DN to DP transition, as T cell-specific deficiency in Zfp335 leads to a substantial accumulation of DN3 along with reduction of DP, CD4^+^ and CD8^+^ thymocytes. This developmental blockade at DN stage results from the impaired intracellular TCRβ expression as well as increased susceptibility to apoptosis in thymocytes. Transcriptomic and ChIP-seq analyses revealed a direct regulation of transcription factors Bcl6 and Rorc by Zfp335. Importantly, enhanced expression of TCRβ and Bcl6/RorγT restores the developmental defect during DN3 to DN4 transition and improves thymocytes survival, respectively. These findings identify a critical role of Zfp335 in controlling T cell development by maintaining intracellular TCRβ expression-mediated β- selection and independently activating cell survival signaling.

## Introduction

T cell development proceeds in a series of developmental stages, which is precisely orchestrated by multiple signaling and molecular networks ^1–3^. Prethymic progenitor cells originated from bone marrow migrate into the thymus and sequentially differentiate into CD4^-^CD8^-^(DN), CD4^+^CD8^+^(DP), and the CD4^+^ or CD8^+^ (SP) stage. Based on the expression of CD44 and CD25, DN thymocytes are divided into several phenotypically distinct stages, including DN1 to DN4 ^4–7^. In the presence of Notch signaling, early thymic progenitor (ETP)-DN1 cells transit into DN2a stage, initiating the T cell lineage commitment, which is immediately accompanied by TCRβ gene arrangement. The majority of DN2 cells enter the DN3 stage with αβ lineage potential ^8^. Only DN3 cells with a complete pre-TCR complex, which consists of the functional TCRβ protein, pre-Tα (pTα) chain and CD3 molecule, can successfully trigger the subsequent maturation into DN4 and DP thymocytes. Further differentiation into mature CD4^+^ or CD8^+^ T cells requires positive and negative selection at DP stage before they migrate to peripheral lymphoid organs ^9–12^.

Pre-TCR signals regulate thymocyte differentiation by mediating protection from apoptosis, stimulating proliferation, and inducing allelic exclusion at the TCRβ locus in post-β-selection DN3b cells and promoting DN to DP transition ^10, 13, 14^. Inactivation of pre-TCR components dampens T lymphocyte development by arresting thymocytes at the DN3 stage and inducing apoptosis ^15–18^. Multiple transcription factors downstream of pre-TCR signaling are involved in T cell differentiation and survival. The major pre-TCR signaling is conducted through the dose-dependent expression of Notch controlled by the Id3-E2A axis ^19^. Abrogation of either Notch or E2A expression may lead to the developmental block of thymocytes at multiple stages ^20–22^. In addition, activation of NF-κB^23^, Ets1^24^ and NFAT5^25^ by pre-TCR signals also contributes to the developmental block. The transcription factor T cell factor 1 (TCF1), together with its downstream Bcl-11b, not only increases the potential to differentiate into T cells ^26, 27^, but also positively regulates thymocyte development via promoting TCRβ recombination and expression, as well as DP cell survival ^28, 29^. Apart from the essential role in the T follicular helper cell lineage commitment ^30^, Bcl6 induced by pre-TCR signals is also involved in the DN to DP transition and protection of DN4 cells from apoptosis ^31^. Additionally, abrogation of RorγT expression in thymocytes leads to a decreased DP proportion and impaired DP survival in a Bcl-xl-dependent manner ^32–34^.

Although pre-TCR signaling is crucial for the β-selection checkpoint, it is not sufficient for progression to the DN3 stage. Other pathways or transcription factors independent of conventional pre-TCR signaling are found to play important roles in the process ^35^ The developmental blockade in Smarca5 or Nkap-deficient thymocytes is pre-TCR-independent, confirmed by intact pre-TCR signals in these mice ^17, 25, 36, 37^. Overall, it remains largely unknown which factors are crucial for T cell development through mechanisms independent of pre-TCR signaling.

Zfp335, also known as the nuclear hormone receptor coregulator (NRC) – interacting Factor 1 (NIF-1), is a zinc finger protein with a 13 C2H2 zinc finger repeating structure consisting of 1337 amino acids^38^. The C2H2-ZF family encodes more than 700 proteins in the human genome, some of which play important roles in ontogenesis, immune cell differentiation, and disease occurrence ^26, 39^, yet the biological characteristics and functions of most members are unclear ^40, 41^. Zfp335 regulates gene transcription by recruiting H3K4 methyltransferase complexes, interacting with co-activators, or directly binding to certain gene promoters ^38, 42–44^. Zfp335 plays important regulatory roles in early embryonic development and neurogenesis ^42^. Germline knockout of Zfp335 is embryonic lethal, while deletion of Zfp335 gene in nerve cells impairs the proliferation and differentiation of nerve progenitor cells in mice, eventually leading to severe microcephaly ^42^. Function of Zfp335 in T cell development has been observed in the study of the Zfp335bloto allele, a missense mutation derived from ENU mutagenesis. While thymocyte development is not significantly affected by this hypomorph mutation, there is a significant reduction in the number of peripheral T cells due to defect in the maturation and migration of thymocytes ^45^. Without loss of function studies, it remains to be determined whether Zfp335 is required for intrathymic T cell development.

In this study, we investigated Zfp335 expression during different thymocyte stages, as well as its function at the β-selection checkpoint and during DN to DP transition. We found that in the thymus, Zfp335 showed a 2- to 10-fold increase in DN3 cells. T cell-specific Zfp335 knockout mice exhibited a profound and remarkable defect in during αβ T cell development along with T cell lymphopenia in peripheral lymphoid organs. Zfp335 deficiency arrested the development of T cell precursors at DN3 stage, accompanied by decreased intracellular TCRβ expression in DN4 cells and increased susceptibility to apoptosis. OT-1 transgenic TCR rescued differentiation from DN3 to DN4 stage, but not at the DP stage. The transcriptomic and ChIP-seq analyses of DN thymocytes revealed that Zfp335 directly targets Bcl6 and Rorc loci. Notably, retroviral overexpression of Bcl6 and Rorc restored DP thymocyte population. Taken together, our findings revealed that Zfp335 support the transition from DN to DP stage by maintaining intracellular TCRβ expression, as well as by promoting DN and DP thymocyte survival via directly regulating Bcl6 and RorγT expression.

## Results

### Impaired thymic αβT cell development in Zfp335-deficient mice

To study the role of Zfp335 in T cell development, we first assessed the expression of Zfp335 among different thymocyte subsets, including DN3, DN4, DP, CD4, CD8, NKT and γδ T cells. We found that DN3 cells displayed relatively high levels of Zfp335 expression (Figure 1A). Consistently, microarray data from ImmGen showed higher expression of Zfp335 specifically at the DN3a stage during T cell development from ETP to CD4/CD8 SP (Figure S1A). Although RNA-seq data from ImmGen exhibited the highest expression of Zfp335 in DP thymocytes, a gradually increased expression was observed from ETP to DN3 (Figure S1B). Given the importance of DN3 stage during β- selection checkpoint, we obtained T cell-specific Zfp335 mice by crossing Zfp335^flox/flox^ strain with Lck-Cre strain (Figure S2A). Zfp335 deletion was confirmed by real-time PCR (qPCR) analysis in DN4 cells (Figure S2B). Strikingly, LckCre^+^Zfp335^f/f^ (KO) mice exhibited significantly smaller thymi and drastically decreased thymocyte numbers than WT control (Figure 1B). Further analysis showed that both percentages and numbers of DP cells, as well as in CD4 SP and CD8 SP cells, were considerably reduced in KO mice (Figure 1C-E). Conversely, the percentage of DN cells was increased by nearly 10-15 folds, although the total number was decreased (Figure 1C, 1D and 1E). In secondary lymphoid organs, we also observed reduced CD4^+^ and CD8^+^ cells in the spleen and lymph nodes (Figure 1F-J). Thus, Zfp335 is essential for the development of αβT cells in the thymus.

**Fig. 1.**
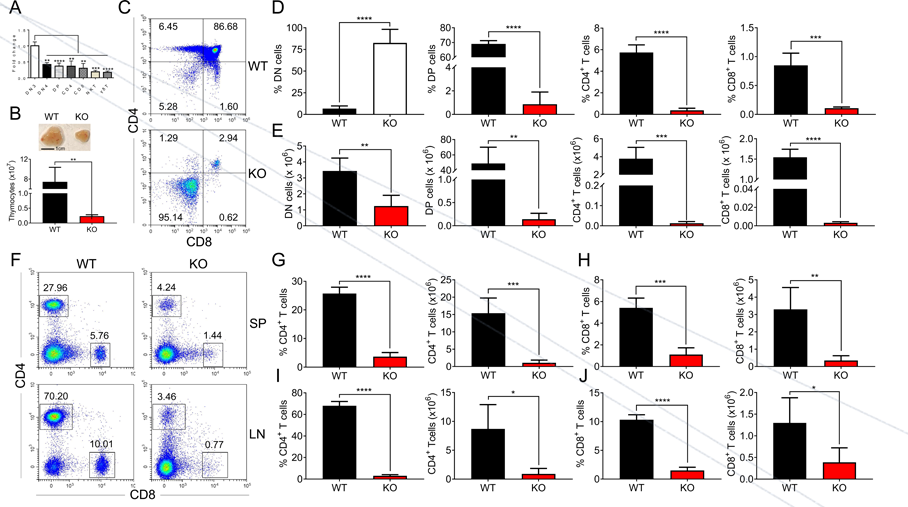
Impaired thymocyte development in Zfp335-deficient mice. **(A)** DN3, DN4, DP, CD4, CD8, NKT and γδT cells were sorted from C57BL/6 thymocytes by flow cytometry. The mRNA levels of Zfp335 were measured by qPCR. **(B)** Thymi from *Lck*Cre^+^*Zfp335*^+/+^ (WT) and *Lck*Cre^+^*Zfp335*^fl/fl^ (KO) mice. Representative thymi and total cell number of thymocytes. Scale bar, 1 cm. **(C-E)** The different stages of thymocyte development in WT and KO mice were measured by flow cytometry. **(C)** Representative FACS plots of DN, DP, CD4, CD8 thymocytes; **(D)** The percentages of DN, DP, CD4, CD8 thymocytes; **(E)** The numbers of DN, DP, CD4, CD8 thymocytes. **(F-J)** CD4^+^ and CD8^+^ cells in spleen and lymph nodes from WT and KO mice were measured by flow cytometry. **(F)** Representative FACS plots of CD4^+^ and CD8^+^ cells in spleen and lymph nodes; **(G-H)** The percentage and number of CD4^+^ T cells **(G)** and CD8^+^ T cells **(H)** in the spleen from WT and KO mice; **(I-J)** The percentage and number of CD4^+^ T cells **(I)** and CD8^+^ T cells **(J)** in the lymph nodes from WT and KO mice. Results represent three independent experiments. n=3 per group. *p < 0.05, **p < 0.01, ***p < 0.001, and ****p < 0.0001.

### Zfp335 intrinsically regulates T cell development in the thymus

To address whether Zfp335 deletion intrinsically affects T cell development, Lin^-^ CD25^+^ CD44^-^ DN3 cells from WT or KO mice were harvested and plated with OP9-DL1 cells in the presence of Flt3L and IL-7, an *in vitr*o model for T-cell development (Figure 2A) ^46^. On both day 2 and day 4, KO group produced fewer DP cells than WT control (Figure 2B-D). DN3 cells from KO mice also generated fewer γδ T cells, despite a higher percentage, in comparison to WT controls after culturing for 4 days (Figure S3A-3C). When DN3 cells from WT (CD45.1^+^) and KO (CD45.2^+^) mice were mixed at a 1:4 ratio, significantly lower percentage of DP cells was also observed in KO group (Figure 2E-G). Furthermore, *in vivo* T cell development was investigated by adoptively transferring T and B cells-depleted bone marrow cells from WT (CD45.1^+^) and KO (CD45.2^+^) mice with 1:4 ratio into WT (CD45.1^+^CD45.2^+^) recipients (Figure 2H). When thymocytes were analyzed 5 weeks post transfer, the percentages of DP, CD4 and CD8 cells were significantly reduced while DN percentage was increased in the mice that received KO cells (Figure 2I-J). Consistently, when WT and KO bone marrow cells were mixed at a 1:4 ratio for co-transfer experiments, fewer DP cells were detected in KO compared to WT group (Figure 2K-L). Together, we demonstrated that Zfp335 intrinsically regulate thymocyte development from DN to DP stage.

**Fig. 2.**
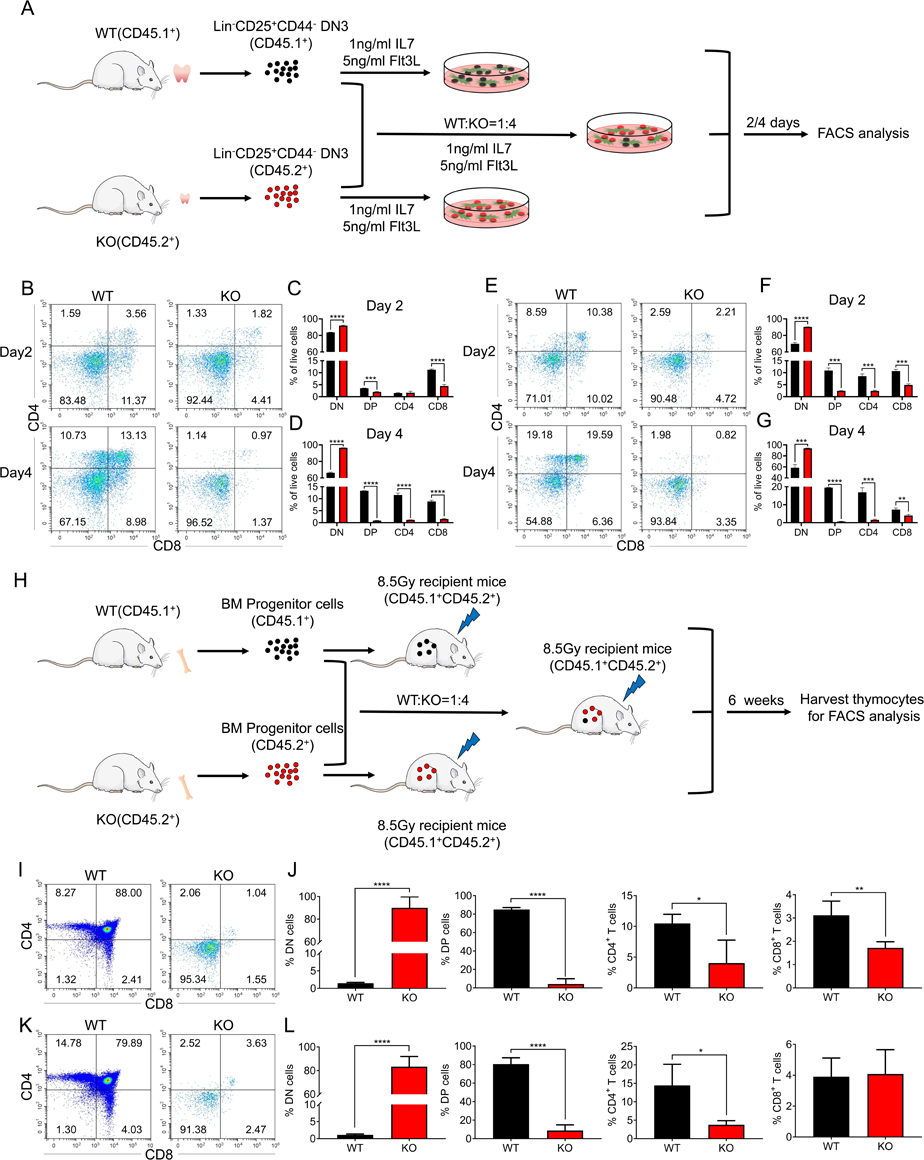
An intrinsic block from DN to DP stage in Zfp335 deficient mice. **(A)** Schematic overview of the *in vitro* OP9-DL1 stromal co-culture assay for T cell differentiation from WT (CD45.1^+^) and KO (CD45.2^+^) DN3 thymocytes to DP and SP thymocytes. **(B-D)** WT and KO DN3 thymocytes were cultured with OP9-DL1 feeder cells *in vitro* in the presence of IL-7 (1ng/ml) and Flt3L (5ng/ml) for 2 and 4 days. The DN and DP thymocytes were measured by flow cytometry. **(B)** Representative FACS plots of DN, DP, CD4^+^ and CD8^+^ thymocytes; **(C)** The percentages of DN, DP, CD4^+^ and CD8^+^ thymocytes 2 days post culture *in vitro*; **(D)** The percentage of DN, DP, CD4^+^ and CD8^+^ thymocytes 4 days post culture *in vitro*. **(E-G)** A mixed population of WT and KO DN3 thymocytes at a 1:4 ratio was co-cultured with OP9-DL1 feeder cells *in vitro* in the presence of IL-7 (1ng/ml) and Flt3L (5ng/ml) for 2 and 4 days. The DN and DP thymocytes were phenotyped by flow cytometry. **(E)** Representative FACS plots of DN, DP, CD4^+^ and CD8^+^ thymocytes; **(F)** Percentages of DN, DP, CD4^+^ and CD8^+^ thymocytes 2 days post culture *in vitro*; **(G)** Percentages of DN, DP, CD4^+^ and CD8^+^ thymocytes 4 days post culture *in vitro*. **(H)** Schematic overview of the *in vivo* bone marrow chimeric mice model for T cell differentiation from WT (CD45.1^+^) and KO (CD45.2^+^) progenitors cells to DP and SP thymocytes. **(I-J)** Full chimeric mice were generated by transplanting WT (CD45.1^+^) or KO (CD45.2^+^) bone marrow progenitor cells into lethally irradiated (8.5 Gy) WT recipient mice (CD45.1^+^CD45.2^+^). Six weeks after transplantation, thymi from recipient mice were harvested. **(I)** Representative FACS plots of DN, DP, CD4^+^ and CD8^+^ thymocytes; **(J)** The percentages of DN, DP, CD4^+^ and CD8^+^ thymocytes. **(K-L)** Full chimeric mice were generated by transplanting a mixed population of WT (CD45.1^+^) and KO (CD45.2^+^) bone marrow progenitor cells at a 1:4 ratio into lethally irradiated WT recipients (CD45.1^+^CD45.2^+^) with 8.5 Gy. Six weeks after transplantation, thymi from recipient mice were harvested. **(K)** Representative FACS plots of DN, DP, CD4^+^ and CD8^+^ thymocytes; **(L)** Percentages of DN, DP, CD4^+^ and CD8^+^ thymocytes. Results represent three independent experiments. n=3 per group. *p < 0.05, **p < 0.01, ***p < 0.001, and ****p < 0.0001.

### Loss of Zfp335 blocks the transition from DN3 to DN4 stage

We further examined the impact of Zfp335 deletion on DN thymocyte development. Staining of CD44 and CD25 on pre-gated lineage (CD4/CD8/TCRβ/TCRδ/NK1.1/CD19/CD11b/CD11c)-negative thymocytes was performed in WT and KO mice. The results showed that a higher percentage of CD44^-^CD25^+^ DN3 cells but a lower percentage of CD44^-^CD25^-^ DN4 cells were found in KO group compared to WT group (Figure 3A and 3B), in which the numbers of both DN3 and DN4 thymocytes were decreased (Figure 3C). The developmental blockade from DN3 to DN4 in Zfp335-deficient cells was verified by *in vitro* co-culture assays on day 2 using OP9-DL1 cultured with WT and KO DN3 cells respectively (Figure 3D-3E) or mixed at a 1:4 ratio (Figure 3F-3G) as described above. The *in vivo* bone marrow chimera models further confirmed the development black at the DN3 stage (Figure 3H-3K), indicating that Zfp335 is indispensable for DN3 to DN4 transition during early stage differentiation.

**Fig. 3.**
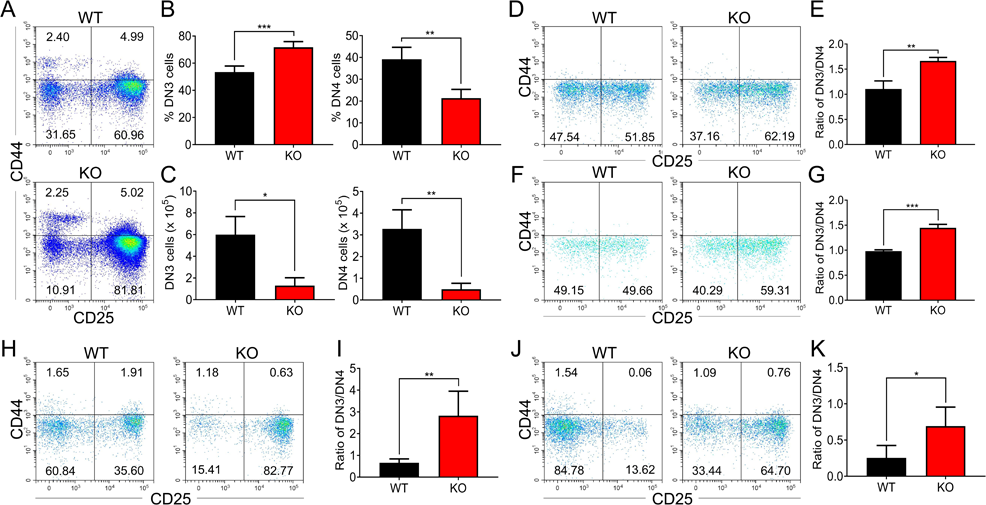
Zfp335-deficient thymocytes undergo a developmental block during DN3 to DN4 transition. **(A-C)** Thymi were harvested from 6- to 8-week-old WT and KO mice. The different stages of DN thymocytes in WT and KO mice were measured by flow cytometry. **(A)** Representative FACS plots of DN1 (CD25^-^CD44^+^), DN2 (CD25^+^CD44^+^), DN3 (CD25^+^CD44^-^), and DN4 (CD25^-^CD44^-^) thymocytes; **(B)** The percentages of DN3 and DN4 thymocytes; **(C)** The numbers of DN3 and DN4 thymocytes. **(D-E)** WT and KO DN3 thymocytes were cultured with OP9- DL1 feeder cells *in vitro* in the presence of IL-7 (1ng/ml) and Flt3L (5ng/ml) for 2 days. The expression of CD44 versus CD25 were measured by flow cytometry. **(D)** Representative FACS plots of DN3 and DN4 thymocytes; **(E)** The ratio of DN3 to DN4 thymocytes 2 days post culture *in vitro*. **(F-G)** A mixed population of WT and KO DN3 thymocytes at a 1:4 ratio was co-cultured with OP9-DL1 feeder cells *in vitro* in the presence of IL-7 (1ng/ml) and Flt3L (5ng/ml) for 2 and 4 days. The expression of CD44 versus CD25 were measured by flow cytometry. **(F)** Representative FACS plots of DN3 and DN4 thymocytes; **(G)** The ratio of DN3 to DN4 thymocytes 2 days post culture *in vitro*. **(H-I)** Full chimeric mice were generated by transplanting WT (CD45.1^+^) or KO (CD45.2^+^) bone marrow progenitor cells into lethally irradiated (8.5 Gy) WT recipients (CD45.1^+^CD45.2^+^). Six weeks after transplantation, thymi from recipient mice were harvested. The expression of CD44 versus CD25 were measured by flow cytometry. **(H)** Representative FACS plots of DN3 and DN4 cells; **(I)** The ratio of DN3 to DN4 thymocytes. **(J-K)** Full chimeric mice were generated by transplanting a mixed population of WT (CD45.1^+^) and KO (CD45.2^+^) bone marrow progenitor cells at a 1:4 ratio into lethally irradiated (8.5 Gy) WT recipient mice (CD45.1^+^CD45.2^+^). Six weeks after transplantation, thymi from recipient mice were harvested. The expression of CD44 versus CD25 were measured by flow cytometry. **(J)** Representative FACS plots of DN3 and DN4 thymocytes; **(K)** The ratio of DN3 to DN4 thymocytes. Results represent three independent experiments. n=3 per group. *p < 0.05, **p < 0.01, and ***p < 0.001.

### Ablation of Zfp335 promotes apoptosis in thymocytes

During the β-selection, efficient proliferation of pre-T cells is necessary for DN to DP progression ^47^, during which the pre-TCR signal functions as a positive regulator of thymocyte survival, allowing for differentiation from pre-T cells into DP thymocytes. We sought to examine whether the defect in Zfp335 KO thymocyte development is due to impaired proliferation or survival. The *in vivo* BrdU incorporation assay showed comparable or even higher percentages of BrdU^+^ DN3 and DN4 cells (Figure 4A-B) and the percentages of Ki67^+^ DN3 and DN4 cells were also similar between the two groups (Figure S4). Nevertheless, when we examined thymocyte apoptosis, KO mice showed a remarkably higher percentage of Annexin V^+^ DN3 and DN4 cells compared to WT cells (Figure 4C-4D). After co-culture with OP9-DL1 for 4 days, both Zfp335-deficient DN3 and DN4 cells showed an increased percentage of Annexin V^+^ cells (Figure 4E-4F). Moreover, in mixed bone marrow chimeras, Zfp335-deficient DN3 and DN4 cells also displayed significantly higher Annexin V^+^ cells (Figure 4G-4H), indicating an intrinsic role of Zfp355 in regulating thymocyte survival.

**Fig. 4.**
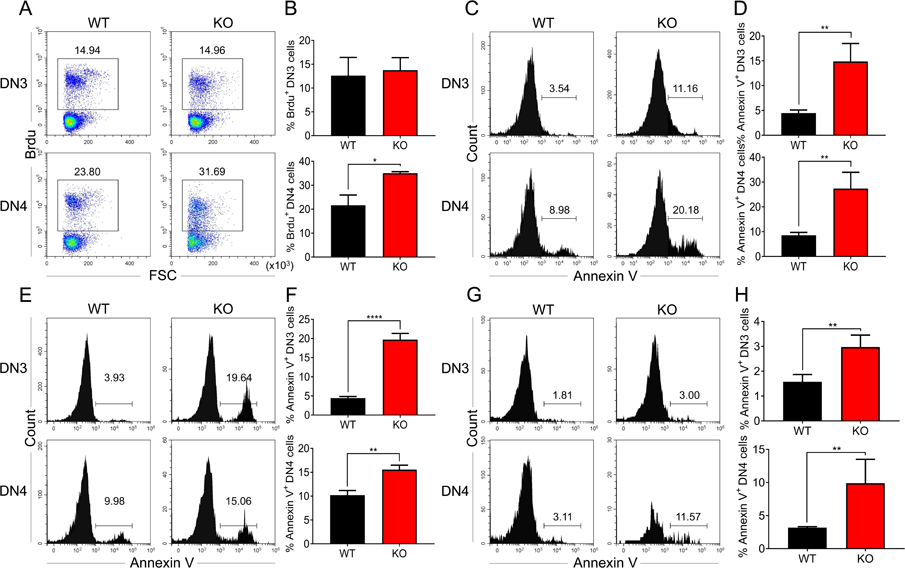
Zfp335 deficiency promoted thymocyte apoptosis *in vivo* and *in vitro*. **(A and B)** Thymi were harvested from 6- to 8-week-old WT and KO mice. The expression of BrdU in DN3 and DN4 thymocytes from WT and KO thymi was measured by flow cytometry. **(A)** Representative FACS plots of BrdU expression in DN3 and DN4 thymocytes; **(B)** The percentages of BrdU^+^ thymocytes in DN3 and DN4 cells. **(C and D)** The binding of Annexin V in DN3 and DN4 thymocytes from WT and KO thymi were measured by flow cytometry. **(C)** Representative FACS plots of Annexin V binding in DN3 and DN4 thymocytes; **(D)** The percentages of Annexin V^+^ thymocytes in DN3 and DN4 cells. **(E-F)** A mixed population of WT and KO DN3 thymocytes at a 1:4 ratio was co-cultured with OP9-DL1 feeder cells *in vitro* in the presence of IL-7 (1ng/ml) and Flt3L (5ng/ml) for 4 days. The binding of Annexin V was measured by flow cytometry. **(E)** Representative FACS plots of Annexin V^+^ DN3 and DN4 thymocytes; **(F)** The percentage of Annexin V binding in DN3 and DN4 thymocytes. **(G-H)** Full chimeric mice were generated by transplanting a mixed population of WT (CD45.1^+^) and KO (CD45.2^+^) bone marrow progenitor cells at a 1:4 ratio into lethally irradiated WT recipient mice (CD45.1^+^CD45.2^+^) with 8.5 Gy. Five weeks after transplantation, thymi from recipient mice were harvested. The binding of Annexin V was measured by flow cytometry. **(G)** Representative FACS plots of Annexin V^+^ DN3 and DN4 cells; **(H)** The percentage of Annexin V binding in DN3 and DN4 thymocytes. Results represent three independent experiments. n=3 per group. *p < 0.05, **p < 0.01, and ****p < 0.0001.

### Ectopic expression of TCRαβ overcomes DN3 stage block in Zfp335- deficient mice

The pre-TCR signal-controlled β-selection is essential for thymocyte differentiation from DN3 to DN4 stage ^48^. We then examined the expression of genes involved in the pre-TCR complex in DN3 and DN4 cell populations. The expression of *Ptcra, Trbc1, Trbc2 and Cd3e* in DN3 and DN4 cells were comparable in WT and KO group (Figure S5A-5B). Additionally, the expression of intracellular CD3 (iCD3) was also unaffected in both Zfp335-deficient DN3 and DN4 cells (Figure S6A-6B). Interestingly, flow cytometry analysis showed no substantial difference in intracellular TCRβ (iTCRβ) expression between WT and KO DN3 cells, whereas Zfp335-deficient mice displayed a significant decrease in the percentage of iTCRβ^+^ DN4 cells (Figure 5A-5B). To further address whether Zfp335 deficiency affects TCRβ expression, we compared differential usage of TCRβ in DN4 cells between WT and KO mice, in which the expression of TCR Vβ5, Vβ6, Vβ8 and Vβ12 were decreased concomitantly (Figure 5C-5F). However, genomic DNA analysis for V-DJβ5 and V-DJβ8 rearrangements showed no difference between WT and KO DN3 cells (Figure S7). Thus, the reduced iTCRβ expression may be a consequence of protein degradation.

**Fig. 5.**
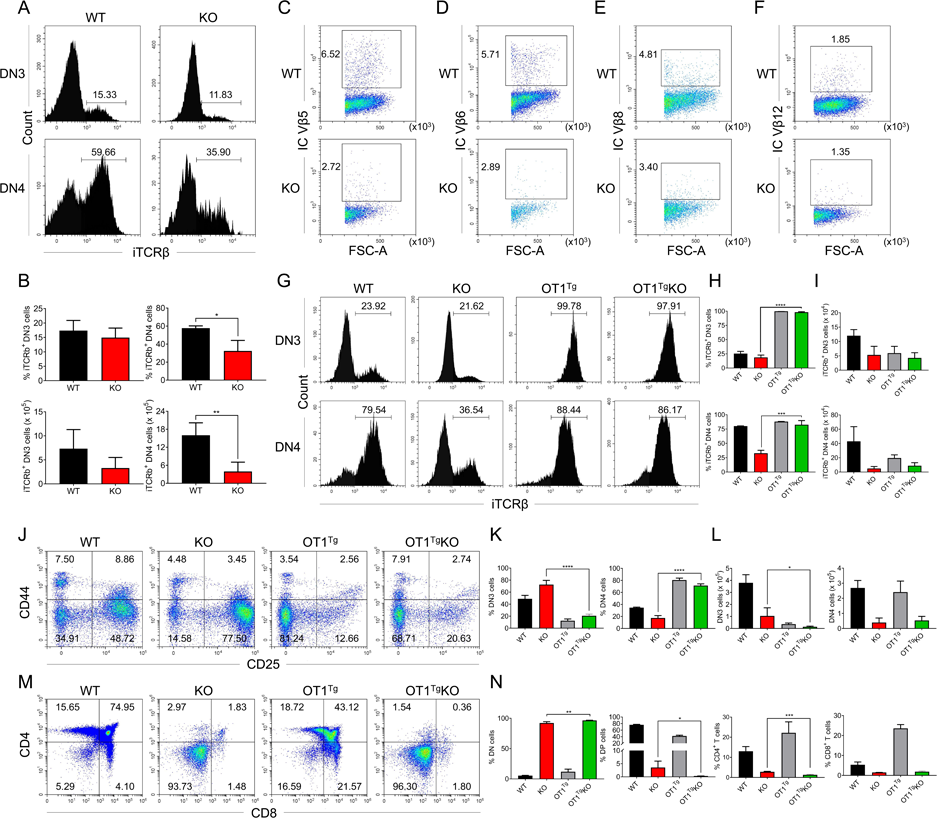
OT1 transgenic TCR overexpression rescued Zfp335-deficiency-induced defect during DN3 to DN4 transition. **(A-B)** Thymi were harvested from 6-8 week-old WT and KO mice. The expression of iTCRβ in DN3 and DN4 thymocytes was measured by flow cytometry. **(A)** Representative FACS plots of iTCRβ expression in DN3 and DN4 thymocytes; **(B)** Percentages and numbers of iTCRβ^+^ thymocytes in DN3 and DN4 cells. **(C-F)** Representative FACS plots of intercellular TCR Vβ5**(C)**, Vβ6**(D)**, Vβ8**(E)**and Vβ12**(F)**expression in DN4 cells from WT and KO mice. **(G-I)** Thymi from WT, KO and *OT1*^+^*Lck*Cre^+^*Zfp335*^fl/fl^ (OT1^Tg^ KO) mice were harvested. The expressions of iTCRβ in DN3 (up) and DN4 (down) thymocytes were measured by flow cytometry. **(G)** Representative FACS plots of iTCRβ expression in DN3 and DN4 thymocytes; **(H)** The percentage of iTCRβ^+^ DN3 and DN4 cells in WT, KO and OT1^Tg^ KO mice; **(I)** The numbers of iTCRβ^+^ DN3 and DN4 cells in WT, KO and OT1^Tg^ KO mice. **(J-L)** The different stages of DN thymocytes in WT, KO, and OT1^Tg^ KO mice were measured by flow cytometry. **(J)** Representative FACS plots of DN3 and DN4 thymocytes; **(K)** The percentages of DN3 and DN4 thymocytes; **(L)** The numbers of DN3 and DN4 thymocytes. **(M-N)** The different stages of thymocyte development in WT, KO, and OT1^Tg^ KO mice were measured by flow cytometry. **(M)** Representative FACS plots of thymocytes. **(N)** The percentages of DN, DP, CD4^+^CD8^-^, and CD4^-^CD8^+^ thymocytes. Results represent three independent experiments. n=3 per group. *p < 0.05, **p < 0.01, ***p < 0.001, and ****p < 0.0001.

Given that Zfp355 deficiency led to diminished iTCRβ expression in DN4 cells, we next investigated the effect of TCR overexpression on aberrant thymocyte development caused by Zfp335 deficiency. Offspring (LckCre^+^Zfp335^f/f^OT1^+^) of LckCre^+^Zfp335^f/f^ mice crossed to OT1 transgenic (Tg) mice was generated to constitutively express TCRα-V2 and TCRβ-V5 Tg gene (OT1^Tg^KO). Notably, forced expression of αβTCR successfully restored the decreased percentage of iTCRβ^+^ DN4 cells in the KO mice, though had little impact on the number of iTCRβ^+^ DN4 cells (Figure 5G-5I). Importantly, developmental arrest at the DN3 stage in Zfp335-deficient mice was fully rescued by OT1 transgene (Figure 5J-5L). However, the proportions of DN, DP, CD4 and CD8 were still comparable in KO and OT1^Tg^KO mice (Figure 5M-5N, Figure S8), demonstrating that the aberrant DP development has not yet been restored by αβTCR overexpression.

### Zfp335 directly targets Bcl6 and Rorc in DN thymocytes

To address the underlying molecular mechanisms of Zfp335-mediated thymocyte development, we performed comprehensive RNA-seq analysis comparing DN4 cells from WT and KO mice. Volcano plot showed that Zfp335- deficient DN4 cells had a total 566 downregulated and 899 upregulated genes (Fold change>1.25, P < 0.05) (Figure 6A and Table 1). Gene ontology (GO) analysis highlighted a large fraction of genes downregulated in KO group belonging to lymphocyte differentiation and apoptotic pathways (Figure 6B). Prominently down-regulated genes associated with lymphocyte differentiation and apoptosis were summarized in the heatmap (Figure 6C). To further determine genes directly regulated by Zfp335, we analyzed Zfp335 by chromatin immunoprecipitation followed by deep sequencing (ChIP-seq). We screened a total of 2797 Zfp335-binding sites (Table 2) and the prevalence of binding peaks across genomic regions was displayed as a venn diagram (Figure S9). When the genome-wide RNA-seq data was combined with Zfp335 ChIP-seq data, 22 genes overlapped between the two datasets (Figure S10A and Table 3). Among these genes, top 10 genes were listed based upon their expression level from RNA-seq result (Figure S10B). qPCR analysis was performed to confirm their downregulation in KO DN4 cells (Figure S10C). Next, we focused on genes related to lymphocyte differentiation and apoptosis. In line with RNA-seq results (Figure 6C), qPCR analysis verified that Bcl6 and Rorc were significantly downregulated in KO DN4 cells (Figure 6D). Importantly, Zfp335 directly targeted promoter regions of Bcl6 and Rorc in ChIP-seq analysis (Figure 6E), which were further verified by luciferase assay (Figure 6F). Taken together, in depth genomic analysis of DN thymocytes supports that Zfp335 directly regulates the transcription of Bcl6 and Rorc.

**Fig. 6.** Zfp335 downstream target analysis in DN4 thymocytes. **(A)** Volcano plot depicting log2 (Fold Change) (x-axis) and −log10 (P value) (y-axis) for differentially expressed genes (abs. log2FC>1, FDR<0.05) in DN4 thymocytes sorted from WT and KO mice; up-regulated (red) and down-regulated (blue). n=2 per group. **(B)** Gene-ontology (GO) analysis of genes that down-regulated in Zfp335-deficient DN4 thymocytes, showing the GO terms related to lymphocyte differentiation and apoptosis (left), the number of genes overlapped with database from the indicated terms (middle left column), P values (middle column) and Q values (middle right column) and genes annotated to the indicated term (right). **(C)** Heatmap of representative genes related to lymphocyte differentiation and apoptosis. The scale ranges from minimum (green boxes) to medium (black boxes) to maximum (red boxes) relative expression. **(D)** The mRNA level of Bcl6 (top) and Rorc (bottom) in DN4 thymocytes from WT and KO mice. **(E)** ChIP-seq analysis for binding of Zfp335 to the Bcl6 and Rorc loci in wild-type DN4 cells. **(F)** Luciferase assay for the binding of different domains of Zfp335 to the promoter regions of Bcl6 and Rorc. The pGL4.16 plasmid was transfected into 293T cells together with MSCV vector carrying different domains. Data represent 3 independent experiments. *p < 0.05, **p < 0.01 and ****p < 0.0001.

### Defect in thymocyte development caused by Zfp335 deficiency can be rescued by Bcl6 and Rorc

To determine whether Bcl6 and Rorc participate in the regulation of thymocyte development downstream of Zfp335, overexpression assays for Bcl6 and Rorc were conducted in *in vitro* thymocyte development model (Figure 7A). DN3 cells from KO mice were co-cultured with OP9-DL1 cells and transduced with retrovirus encoding Mock-GFP, Zfp335-GFP, Bcl6-GFP or Rorc-GFP. After 3.5 days, overexpression of Bcl6 and Rorc resulted in substantial DP generation, particularly in the Bcl6 group, resulting in a similar DP proportion to that in Zfp335-overexpressing cells (Figure 7B-7C). Of note, overexpression of Psmg2, Dctn1, Ankle2, Cep76, Fgf13 and Ddx31 identified in the qPCR results (Figure S10B and S10C) in KO DN3 cells did not restore DP generation *in vitro* (Figure S11). Importantly, enhanced expression of Bcl6 in DN3 cells rescued DN thymocyte apoptosis (Figure 7D-7E), while overexpression of both Bcl6 and Rorc rescued DP thymocytes from enhanced apoptosis (Figure 7F-7G). p53, negatively regulated by Bcl6, is involved in lymphocyte apoptosis ^49–51^. Thus, we crossed Zfp335 KO strain with Trp53 KO strain to obtain double knockout mice, in which Trp53 deletion resulted in a partial recovery of DP percentage, but not cell number (Figure S12), further supporting the role of Bcl6 in Zfp335- controlled thymocyte survival. Together, these data demonstrated that Zfp335 controls DN thymocyte survival through direct regulation of Bcl6 and Rorc expression.

**Fig. 7.** Identification of Bcl6 and Rorc as functional targets of Zfp335 for regulating thymocyte development. **(A)** Schematic overview of the *in vitro* gene overexpression in KO DN3 thymocytes and following T cell differentiation in OP9-DL1 co-culture system. **(B-C)** Zfp335-deficienct DN3 thymocytes (KO) were cultured with OP9-DL1 feeder cells *in vitro*, then transduced with either mock, Zfp335-overexpressing, Bcl6-overexpressing, or Rorc-overexpressing vector for 3.5 days. The different stages of thymocyte development from GFP positive cells were measured by flow cytometry. **(B)** Representative FACS plots of DN and DP thymocytes from the indicated groups; **(C)** The percentage of DP thymocytes from GFP positive cells. **(D-G)** KO DN3 thymocytes were cultured with OP9-DL1 feeder cells *in vitro*, then transduced with either mock, Zfp335-overexpressing, Bcl6- overexpressing, or Rorc-overexpressing vector for 3.5 days. The expressions of Annexin V in DN and DP thymocytes were measured by flow cytometry. **(D)** Representative FACS plots of Annexin V^+^ DN thymocytes from the indicated groups; **(E)** The percentage of Annexin V^+^ DN thymocytes from GFP positive cells; **(F)** Representative FACS plots of Annexin V^+^ DP thymocytes from the indicated groups; **(G)** The percentage of Annexin V^+^ DP thymocytes from GFP positive cells. Results shown represent three independent experiments. *p < 0.05, **p < 0.01, **p < 0.001 and ***p < 0.001.

## Discussion

In this study, we revealed that Zfp335 is essential for thymocyte development, particularly during DN to DP transition. Zfp335 deficiency in T cells led to a significant loss of DP, CD4 SP and CD8 SP cells and an accumulation of DN3 cells. Mechanistically, the developmental blockade is attributed to both impaired pre-TCR signal and increased susceptibility to apoptosis. Serving as a transcription factor, Zfp335 directly promotes Bcl6 and Rorc expression in DN thymocytes to ensure their survival during early development.

We have shown that Zfp335 expression was upregulated specifically in DN3 thymocytes and significantly decreased in the subsequent stages, suggesting a critical role at the DN3 stage. Of note, loss of Zfp335 led to a dramatic reduction in thymus size and thymocyte number. The accumulation of DN3 cells results from an intrinsic mechanism that hinders the transition from DN3 to DN4 stage, leading to the reduction of DP thymocytes and mature T cells in the periphery. These data are in line with another recent study reporting that Zfp335 mutation led to a reduction in peripheral T cells as a result of defective naïve T cells and SP thymocytes ^45^. However, given the intact thymic selection with Zfp335 mutation, the report was inconsistent with our observation of decreased β-selection with Zfp335 deficiency. The discrepancy is likely due to the different approaches used to disrupt Zfp335 function since a single-nucleotide missense mutation of Zfp335 may affect its function differently. Nevertheless, by knocking out the entire Zfp335 protein, we provide evidence that Zfp335 is indispensable for early thymocyte development.

Thymocyte β-selection is a critical developmental checkpoint allowing for the progression from DN3 to DN4 stage and the maintenance of DP cell numbers, which is primarily dependent on TCRβ and pre-TCR signals that require a functional iTCRβ paired with a pre-Tα chain ^52^. Pre-TCR signaling regulates thymocytes differentiation, proliferation, and survival in the full developmental process ^53^. While Zfp335-deficient DN4 cells exhibited no defects in the rearrangement of TCRβ chain genes and pTα gene expression, our results clearly demonstrated that Zfp335 deficiency markedly impaired intracellular TCRβ expression and led to an unbiased reduction of the majority of Vβ genes in DN4 populations. Future studies will investigate the mechanism for how Zfp335 regulates iTCRβ expression. Importantly, forced iTCRβ expression in DN3 and DN4 cells by transduction of OT1-TCR completely rescued the developmental impairment during the DN3-DN4 transition in Zfp335-deficient mice despite the failure to rescue the DN3, DN4 and DP population size. This suggests that Zfp335 controls the DN3-DN4 transition dependent on pre-TCR signals, but other mechanisms may also regulate the DP population size.

The large population of DP thymocytes is maintained by both cell proliferation and survival mechanisms. Zfp335-deficient DN3 and DN4 cells showed slightly higher or unchanged incorporation of BrdU, suggesting that cell proliferation was not affected. However, our data revealed a significant increase in apoptosis in Zfp335-deficient DN3 and DN4 thymocytes. Transcriptomic analysis (RNA-Seq and qPCR) unveiled the downregulation of Bcl6 and RorγT signaling which are critically involved in thymocyte apoptosis ^31, 34^. Indeed, we have demonstrated that Zfp335, a transcription factor, direct binds to the promoter regions of Bcl6 and RorγT genes. More importantly, enhanced expression of Bcl6 and RorγT could improve thymocyte survival and substantially restore the DP thymocyte population. Trp53 deletion resulted in a partial recovery of DP cells, further supporting the role of Bcl6 in Zfp335-controlled thymocyte survival. Of note, Zfp335 may also control thymocyte survival through directly regulating other targets.

In conclusion, our study revealed that the C2H2 zinc finger protein Zfp335 plays a novel and crucial role during thymocyte development, specifically during the transition from DN to DP stage. We further demonstrated that Zfp335 promotes Bcl6 and RorγT signaling to prevent thymocytes apoptosis and ensure the survival and differentiation of thymocytes. Collectively, we provided evidence that Zfp335 is essential for thymocyte development through pre-TCR dependent and independent mechanisms.

## Acknowledgements

We thank Drs. Xiaofei Wang and Guohua Zhang for Flow cytometry analysis and cell sorting. We also thank the Mouse Facility of Xi’an Jiaotong University. This work was supported by the National Natural Science Foundation of China grants 3217080356 (to B.Z.), National Natural Science Foundation of China grants 81771673(to B.Z.), Major International (Regional) Joint Research Project grants 81820108017 (to B.Z.) and the Natural Science Foundation of Shaanxi Province (2020JQ-098, L.L.).

## Contributions

B.Z. designed research and interpreted data. X.W., A.J., Y.S., W.L., R.D., B.Y., X.Y., C.S., L.S., C.Z. and H.L. performed the experiments and analyzed data.

X.W. and C.Z. analyzed the data. B.Z., X.W. and L.S. wrote the paper.

## Materials and Methods

### Mice

*Zfp335*^fl/fl^, *Lck*Cre, and *OT-1* strains were purchased from The Jackson Laboratory (Bar Harbor, ME, USA). *Lck*Cre mice were crossed with *Zfp335*^fl/fl^ mice to generate *Lck*Cre^+^*Zfp335*^fl/fl^ (KO) mice and *Lck*Cre^+^*Zfp335*^+/+^ (WT) mice. Mice aged 6-8 weeks were used for analyses in the study. All mice were housed in specific-pathogen free conditions by the Xi’an Jiaotong University Division of Laboratory Animal Research. All animal procedures were approved by the Institutional Animal Care and Use Committee of Xi’an Jiaotong University.

### Antibodies and reagents

The following antibodies and kits were purchased from Biolegend (San Diego, CA, USA): APC/Cy7 anti-CD4 (clone GK1.5), PE anti-CD8α (clone 53-6.7), PE/Cy7 anti-TCRβ (clone H57-597), FITC anti-TCRγδ (clone GL3), PE/Cy7 anti-CD44 (clone IM7), PE anti-CD25 (clone PC61), FITC anti-NK1.1 (clone PK136), FITC anti-CD19 (clone 6D5), FITC anti-CD11b (clone M1/70), FITC anti-CD11c (clone N418), FITC anti-TER-119/Erythroid Cells (TER-119), APC/Cy7 anti-CD45.1 (clone A20), PE anti-CD45.2 (clone 104), Pacific Blue™ anti-Annexin V (Cat # 640918), Pacific Blue™ anti-CD3 (clone 17A2), PE anti-BrdU (clone Bu20a) and the Fixation/Permeabilization Solution Kit (Cat # 554722). PE anti-Ki67 monoclonal antibody (clone SolA15) was purchased from eBioscience (San Diego, CA, USA). Quick-RNA Microprep Kit (Cat # R1051) was obtained from Zymo Research (Irvine, CA, USA).

### FACS analysis and sorting

Lymphocytes from *Lck*Cre^+^*Zfp335*^+/+^ and *Lck*Cre^+^*Zfp335*^fl/fl^ mice were isolated. For surface staining, single cell suspension was prepared. A total of 1×10^6^ cells were stained in the dark at 4 ℃ for 30 min with indicated antibodies. The analysis was performed on a CytoFLEX flow cytometer (Beckman Coulter; Brea, CA, USA). DN3 (Lin^-^CD25^+^CD44^-^) and DN4 (Lin^-^CD25^-^CD44^-^) cells were collected by BD FACSAria™ Ⅱ cell sorter (BD Biosciences, San Jose, CA, USA). FACS data were recorded and analysed using CytExpert software (Version 2.3.0.84; Beckman Coulter; Indianapolis, IN, USA).

### Intracellular staining

DN and DP thymocytes were phenotyped using a combination of surface antibodies against lineage markers (CD4, CD8α, TCRβ, TCRγδ, NK1.1, CD19, CD11b, CD11c, and Ter119), together with CD44 and CD25 antibodies. For intracellular cytokine staining, the thymocytes were fixed and permeabilized using a Fixation/Permeabilization Solution Kit (Biolegend), followed by staining using indicated antibodies. The cells were analyzed on a CytoFLEX flow cytometer (Beckman Coulter).

### Quantitative RT-PCR

Cell lysis was performed with RNA extraction and cDNA synthesis using Quick-RNA Microprep Kit (Zymo Research) and ReverTra Ace qPCR RT Master Mix Kit (TOYOBO), respectively. The qRT-PCR reactions were carried out using StepOnePlus™ Real-Time PCR Systems (ABI) with SYBR mixture (Genstar) to determine relative gene expression. The sequences for the primers are list in the table 4.

### Bone marrow transplantation

Lineage-negative BM cells from CD45.1^+^ mice and *Lck*Cre^+^*Zfp335*^fl/fl^ mice (CD45.2^+^) were sorted, and mixed at a 1:4 ratio and co-transferred into lethally irradiated (8.5Gy) recipient mice (CD45.1^+^CD45.2^+^). Six weeks after bone marrow transplantation (BMT), thymocytes from recipient mice were harvested for FACS analysis.

### In vitro OP9-DL1 Cell Co-culture

Both Lin^-^CD25^+^CD44^-^ DN3 cells and Lin^-^CD25^-^CD44^-^ DN4 cells were sorted from the thymi of *Lck*Cre^+^*Zfp335*^+/+^ and *Lck*Cre^+^*Zfp335*^fl/fl^ mice and co-cultured with OP9-DL1 feeder cells in α-MEM medium in the presence of IL-7 (1 ng/ml, PeproTech) and Flt3-L (5 ng/ml, PeproTech). On day 2 and day 4, total cells were collected and stained with indicated antibodies for FACS analysis.

### Retroviral transduction of DN3 thymocytes

Retroviruses were produced from BOSC cells transfected with Mock-GFP, Zfp335-GFP, Bcl6-GFP and Rorc-GFP retroviral plasmids. For retroviral transduction, Lin^-^CD25^+^CD44^-^ DN3 thymocytes were sorted by FACSAria flow cytometry (BD) and co-cultured with OP9-DL1 feeder cells overnight in the presence of 1ng/ml IL-7 and 5ng/ml Flt3L. Retroviral infection was performed 16h later by centrifugation (2500 rpm for 90 min at 37°C) in the presence of retroviral supernatants and 8ug/ml polybrene. After spinning, supernatants were replaced by α-MEM medium with 10% FCS supplemented with 1ng/ml IL-7 and 5ng/ml Flt3L. 3.5 days later, GFP^+^ cells were examined using flow cytometry analysis.

### Luciferase Assay

To assess whether Zfp335 regulates Bcl6 and Rorc by directly binding to their promoter regions, the DNA fragments were cloned into the pGL4.16 (luc2CP/Hgro) vector (Promega) which contains the luciferase reporter gene luc2CP. The pGL4.16 plasmid, control vector pGL4.74 (hRluc/TK) encoding the luciferase reporter gene hRluc (Renilla reniformis), along with plasmids expressing candidate genes were transfected separately into 293T cell line (ATCC). 48 hours post-transfection, the luciferase activity of both Firefly and Renilla luciferase was measured using a Dual-Luciferase Reporter kit (Promega) on SYNERGY Neo2 multi-mode reader (BioTek).

### RNA-Seq library preparation and sequencing

Lin^-^CD25^-^CD44^-^ DN4 cells were sorted from the thymi of *Lck*Cre^+^*Zfp335*^+/+^ and *Lck*Cre^+^*Zfp335*^fl/fl^ mice. RNA isolation was performed using the RNeasy Mini Kit (Qiagen) according to the manufacturer’s protocol. RNA quality and quantity were detected by the Qubit RNA broad range assay in the Qubit Fluorometer (Invitrogen). After quality control using RNase-free agarose gel and Agilent 2100 (Agilent Technologies, Palo Alto, CA, USA), RNA-Seq libraries were prepared by using 200ng total RNA with TruSeq RNA sample prep kit (Illumina). Oligo(dT)-enriched mRNAs were fragmented randomly with fragmentation buffer, followed by first- and second-strand cDNA synthesis. After a series of terminal repair, the double-stranded cDNA library was obtained through PCR enrichment and size selection. cDNA libraries were sequenced with the Illumina Hiseq 2000 sequencer (Illumina HiSeq 2000 v4 Single-Read 50 bp) after pooling according to its expected data volume and effective concentration.

Two biological replicates were performed in the RNA-Seq analysis. Raw reads were then aligned to the mouse genome (GRCm38) using Tophat2 RNA-Seq alignment software, and unique reads were retained to quantify gene expression counts from Tophat2 alignment files. Data were analyzed and preprocessed in the R environment. Differential expression analysis was performed using R package DESeq2 (Adjusted P value < 0.05 and fold-change > 1.25). Heat maps and volcano plots were visualized using the R package.

### ChIP-Seq library preparation and sequencing

Both Lin^-^CD25^+^CD44^-^ DN3 cells and Lin^-^CD25^-^CD44^-^ DN4 cells were sorted from the thymi of *WT* mice by FACSAria flow cytometry (BD). Anti-Zfp335 antibody (Novus) and Millipore 17–10085 ChIP kit were used in the ChIP assay. Immunoprecipitated DNA was used for Illumina ChIP-Seq sample preparation. In brief, 5×10^7^ cells were crosslinked to chromatin with 1% formaldehyde. Reaction was stopped with 0.125M glycine. Cells were then resuspended in cold nuclear lysis buffer and sonicated to obtain DNA with ∼300-500bp size, followed by precipitation by incubation with immunoprecipitation–grade anti- Zfp335 antibody and Magnetic Protein A/G Beads overnight. The following day, beads were sequentially washed by low-salt, high-salt, LiCl, and TE buffers. Bound complexes were eluted in 150μl of elution buffer at 62°C for 2h with shaking, followed by reversal of formaldehyde crosslinking at 95°C for 10 minutes. DNA was eventually purified with spin columns.

The concentration of immunoprecipitated DNA was detected by the Qubit DNA broad range assay in the Qubit Fluorometer (Invitrogen). 10ng immunoprecipitated DNA was prepared for sequencing using the Illumina ChIP-Seq sample preparation protocol. Blunt-end DNA fragments were ligated to Illumina adaptors, amplified, and sequenced using the SE150 model. Raw reads were filtered firstly to remove low-quality or adaptor sequences by SOAPnuke (version 1.5.6). Clean reads were mapped to the reference genome of GRCm39 with SOAPaligner/soap2 (version 2.21t) using default settings. The MACS2 software (Version 2.1.1) was used to process peak calling. The different enrichment peaks from different samples were plotted by MAnorm (version 1.1). Genomic graphs were generated and viewed with the IGV (Integrative Genomics Viewer).

### Statistical analysis

Data were presented as mean ± SEM. Statistical analyses were applied to biologically independent mice or technical replicates for each experiment which was independently repeated at least three times. Two-tailed Student’s t-test was used for all statistical calculations using GraphPad Prism 7 software. All bar graphs include means with error bars to show the distribution of the data. The level of significance is indicated as follows: *P < 0.05, **P < 0.01, ***P < 0.001, ****P < 0.0001.

## Data availability

The sequencing data presented in this paper are available for download on GEO data repository with accession numbers GEO: GSE184532 and GSE184705.

## Supplemental Figure legend

**Fig. S1.**
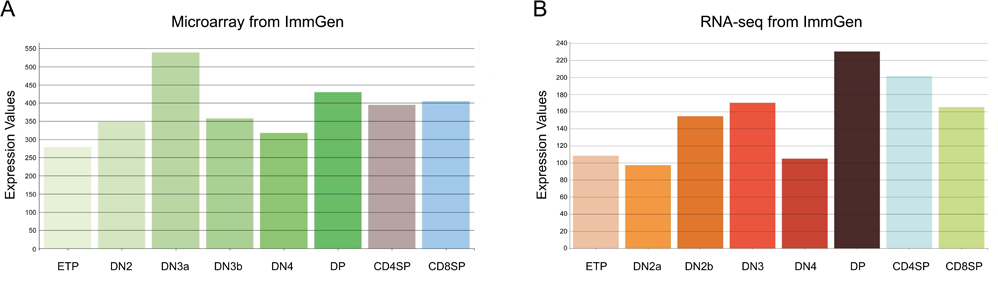
The transcriptional profiling of Zfp335 expression in various subsets of thymocytes, related to Fig. 1. **(A-B)** The RNA profiling data sets of Zfp335 generated by microarray **(A)** or RNA-seq **(B)** data on the ImmGen database.

**Fig. S2.**
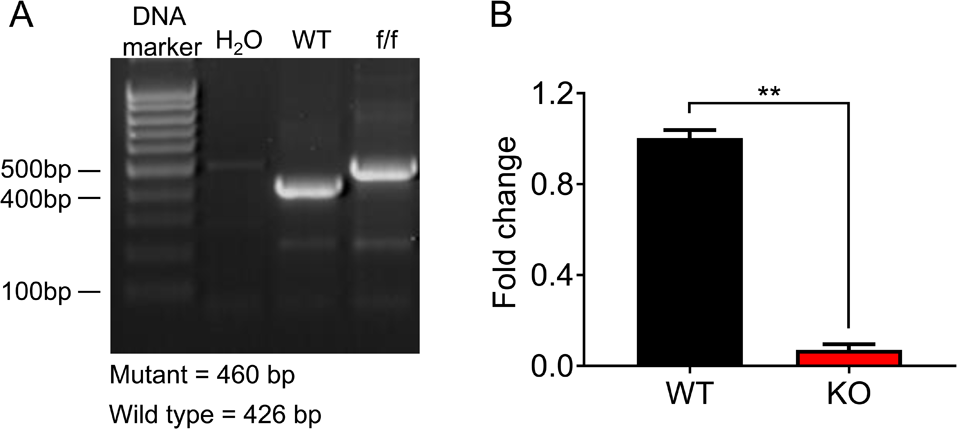
Verification of Zfp335 conditional knock-out mouse strain, related to Fig. 1. **(A-B)** Zfp335^f/f^ mice were crossed with LckCre mice to generate LckCre^+^Zfp335^f/f^ mice. **(A)** The size of Zfp335 DNA in WT and KO mice; **(B)** The mRNA level of Zfp335 in DN4 cells from WT and KO mice. **P < 0.01, n=3.

**Fig. S3.**
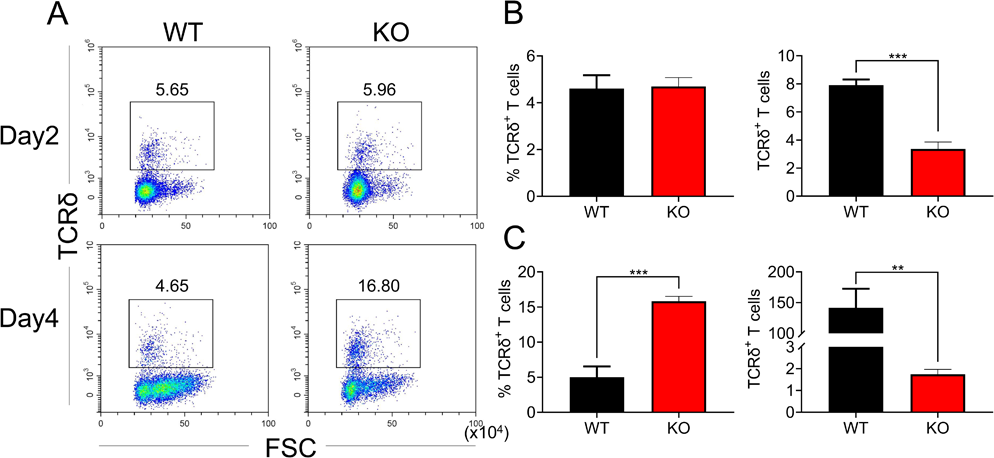
Effect of Zfp335 deletion on γδ T cell percentages and cellularity in vitro, related to Fig. 1. WT and KO DN3 thymocytes were cultured with OP9-DL1 feeder cells *in vitro* in the presence of IL-7 (1ng/ml) and Flt3L (5ng/ml) for 2 and 4 days. The γδ T cells were measured by flow cytometry. **(A)** Representative FACS plots of γδ T cells; **(B and C)** The percentages of γδ T cells on day2 **(B)** and day4 **(C)**. Data are representative of at least three independent experiments shown as the mean ± s.e.m. Two-tailed, unpaired t-tests were used for statistical analyses;**P ≤ 0.01, ***P ≤ 0.001.

**Fig. S4.**
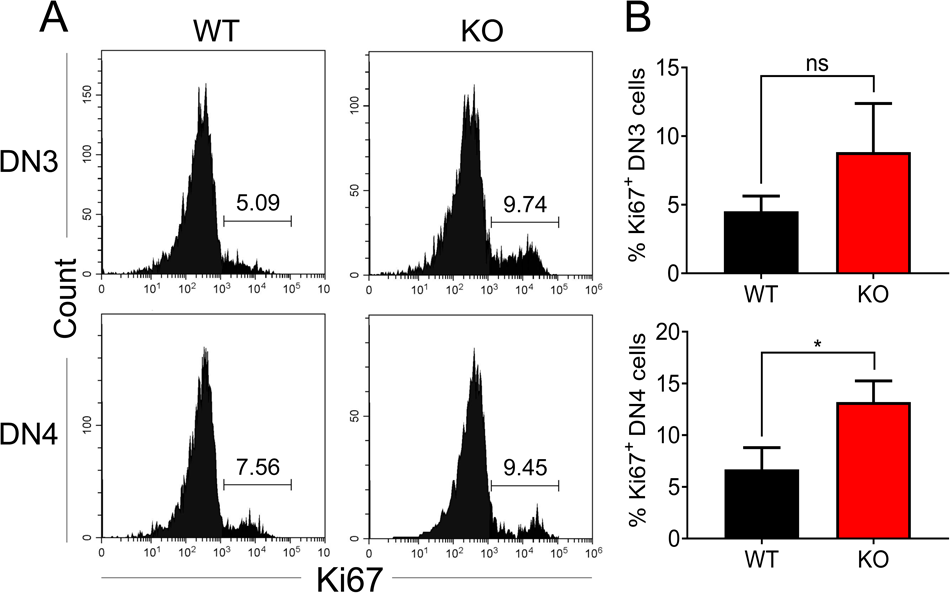
Zfp335 deficiency have no effect on the proliferation of DN3 and DN4 cells. **(A and B)** Thymi from WT and KO mice were harvested. The expressions of Ki67 in DN3 and DN4 thymocytes from WT and KO thymi were measured by flow cytometry. **(A)** Representative FACS plots of Ki67 expression in DN3 and DN4 thymocytes; **(B)** The percentages of Ki67^+^ thymocytes in DN3 and DN4 cells. Data are representative of at least three independent experiments shown as the mean ± s.e.m. Two-tailed, unpaired t-tests were used for statistical analyses; *p < 0.05.

**Fig. S5.**
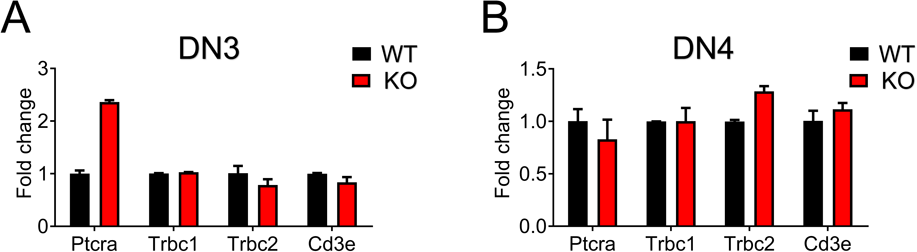
Effect of Zfp335 deletion on pre-TCR complex expression in DN3 and DN4 Cells. qPCR analysis of Ptcrα, Trbc1,Trbc2 and CD3e expression in DN3 **(A)** and DN4 **(B)** cells from WT and KO mice.

**Fig. S6.**
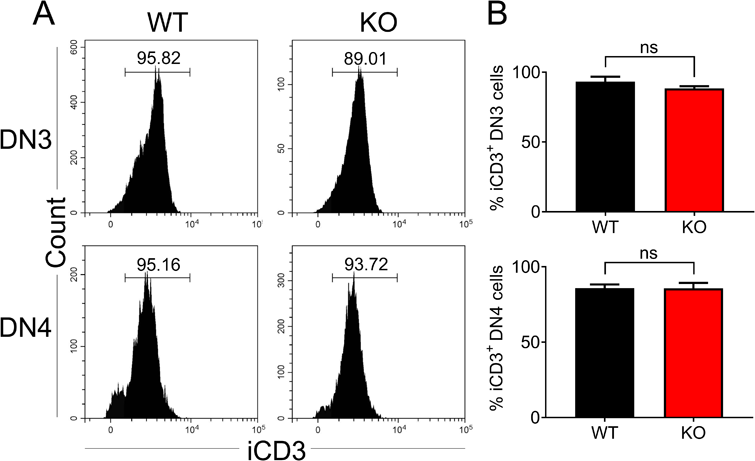
Intercellular CD3 expression in Zfp335-deficienct DN3 and DN4 thymocytes. **(A)** Representative FACS plots of intercellular CD3 (iCD3) expression in DN3 and DN4 thymocytes. **(B)** The percentages of iCD3^+^ DN3 and DN4 cells in WT and KO mice. Data are representative of at least three independent experiments shown as the mean ± s.e.m. Two-tailed, unpaired t-tests were used for statistical analyses.

**Fig. S7.**
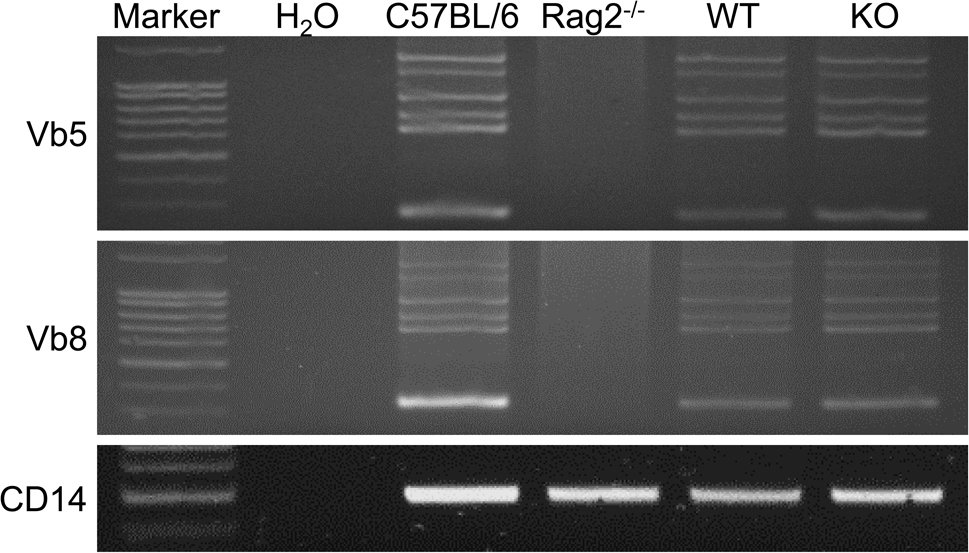
Effect of Zfp335 deletion on TCRβ rearrangement in DN3 thymocytes. PCR analysis of TCRβ gene rearrangements. Genomic DNA isolated from thymocytes of B6, recombinase-activating gene-2-deficient (*Rag2*^−/−^), WT, and KO mice was amplified with primers that detect rearrangements both Vβ5 and Vβ8. Data are representative of three replicates.

**Fig. S8.**
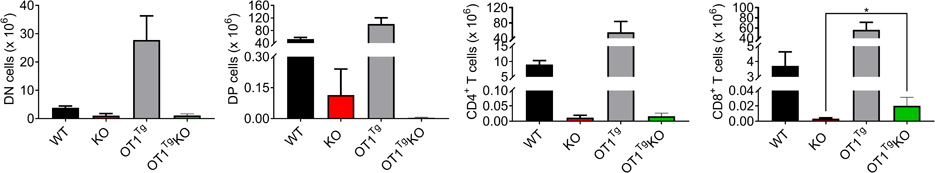
OT1Tg failed to rescue thymocyte numbers. The numbers of DN, DP, CD4^+^CD8^-^, and CD4^-^CD8^+^ thymocytes in WT, KO, and OT1^Tg^ KO mice were measured.

**Fig. S9.**
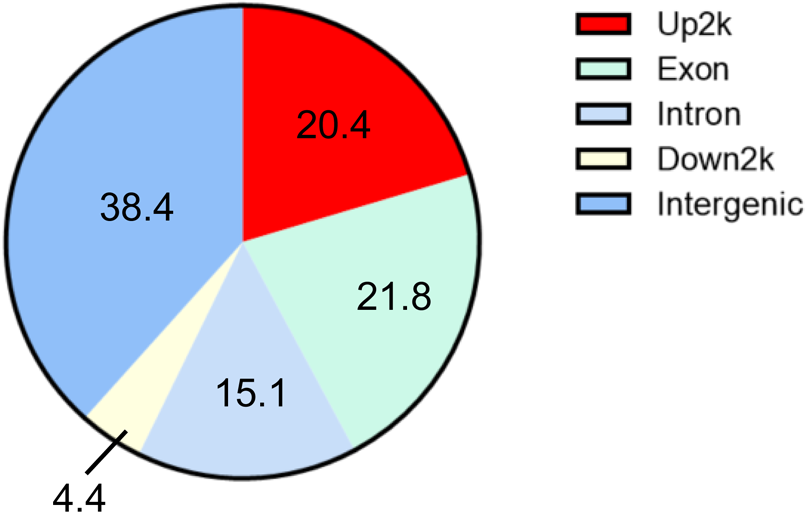
ChIP-Seq analysis of Zfp335 binding sites. The distribution of peaks in various regions of the genome was calculated.

**Fig. S10.**
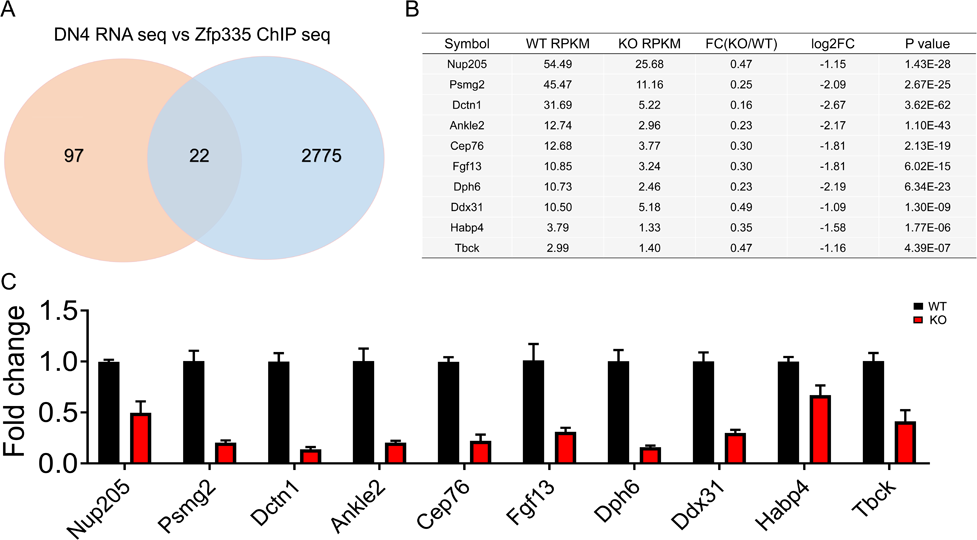
The intersection analysis of Zfp335 by RNA-Seq and ChIP-Seq. **(A)** Venn diagram depicting the overlapping portion of differentially expressed genes in RNA-Seq (red) and Zfp335 binding regions in ChIP-Seq (blue); **(B)** The shared top 10 overlapped genes from the RNA-Seq and ChIP-Seq; **(C)** qPCR analysis of the top 10 overlapped genes depicted in **(B)**.

**Fig. S11.**
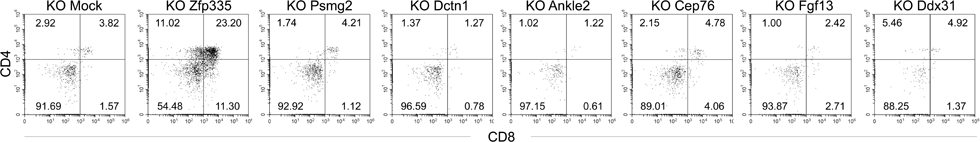
Identification of the top 10 overlapped genes in KO DN3 thymocytes for the regulation of thymocyte development. Zfp335 deficiency DN3 thymocytes (KO) were cultured with OP9-DL1 feeder cells *in vitro*, then transduced with the top 10 overlapped genes overexpression vector for 3.5 days. The different stages of thymocyte development were measured by flow cytometry.

**Fig. S12.**
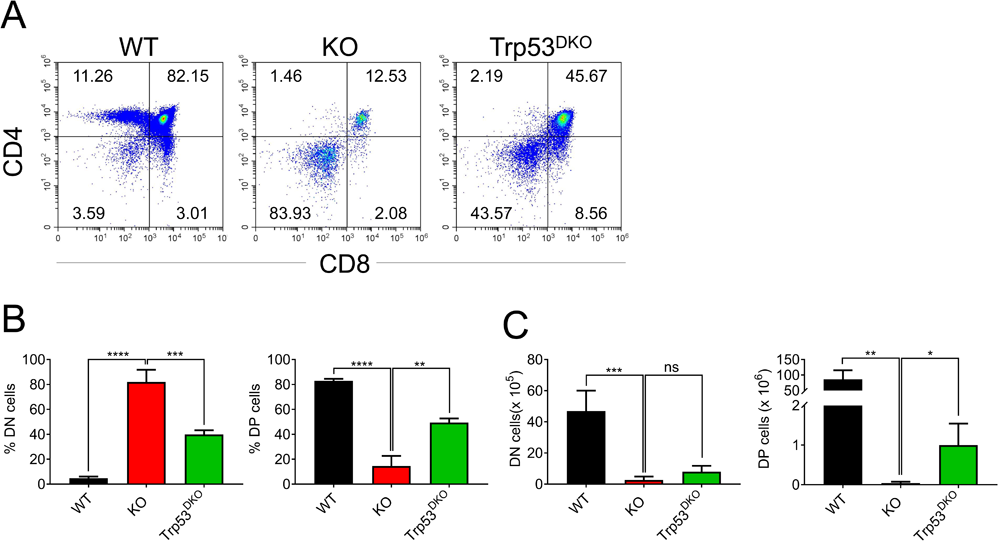
Examination of Trp53 as the target of Zfp335 during the regulation of thymocyte development. Thymi from WT, KO, and *Trp53*^KO^*Lck*Cre^+^*Zfp335*^fl/fl^ (Trp53^DKO^) mice were harvested. The different stages of thymocyte development in WT, KO, and Trp53^DKO^ mice were measured by flow cytometry. **(A)** Representative FACS plots of thymocytes. **(B)** The percentages of DN, DP, CD4^+^CD8^-^, and CD4^-^CD8^+^ thymocytes. **(C)** The numbers of DN, DP, CD4^+^CD8^-^, and CD4^-^CD8^+^ thymocytes. Data are representative of at least three independent experiments shown as the mean ± s.e.m. Two-tailed, unpaired t-tests were used for statistical analyses; *p < 0.05, **p < 0.01, ***p < 0.001, and ****p < 0.0001.

